# Multimodal characterization of Yucatan minipig behavior and physiology through maturation

**DOI:** 10.1101/2021.03.18.436053

**Authors:** Alesa H. Netzley, Ryan D. Hunt, Josue Franco-Arellano, Nicole Arnold, Kirk A. Munoz, Aimee C. Colbath, Tamara Reid Bush, Galit Pelled

## Abstract

Brain injuries that are induced by external forces are particularly challenging to model experimentally. In recent decades, the domestic pig has been gaining popularity as a highly relevant animal model to address the pathophysiological mechanisms and the biomechanics associated with head injuries. Understanding cognitive, motor, and sensory aspects of pig behavior throughout development is crucial for evaluating cognitive and motor deficits after injury. We have developed a comprehensive battery of tests to characterize the behavior and physiological function of the Yucatan minipig throughout maturation. Behavioral testing included assessments of learning and memory, executive functions, circadian rhythms, gait analysis, and level of motor activity. We applied traditional behavioral apparatus and analysis methods, as well as state-of-the-art sensor technologies to report on motion and activity, and artificial intelligent approaches to analyze behavior. We studied pigs from 16 weeks old through sexual maturity at 35 weeks old. The results show multidimensional characterization of minipig behavior, and how it develops and changes with age. This animal model may capitulate the biomechanical consideration and phenotype of head injuries in the developing brain and can drive forward the field of understanding pathophysiological mechanisms and developing new therapies to accelerate recovery in children who have suffered head trauma.

## Introduction

Brain disorders, diseases and injuries caused by external forces, such as sports collisions, car accidents, falls, violent attacks and blasts, remain challenging to model experimentally (1–3). The nature and the extent of the injury depends upon a complex array of anatomical, physiological, and biomechanical parameters, including the size of the brain, thickness of skull, the ratio of grey to white brain matter, age at the time of injury, as well as the energy, angle, and acceleration of the impact itself (1, 4, 5). A single event or series of events may result in changes in cognition, as well as sensorimotor and physiological impairments that are often difficult to measure in experimental models (6–8).

An emerging animal model for brain injury research is the domestic pig (*sus scrofa domesticus*). There are distinct advantages to using a large animal model for *in vivo* brain injury research (9–12). Compared to rodent models, which are the most commonly used animal models for central nervous system injury studies, pigs are much more similar to humans in terms of brain size (13), neuroanatomy (13), brain organization and morphology (14, 15), neurodevelopment (7, 16, 17), and neuroinflammatory mechanisms (18–21). Due to the strong similarities between pigs and humans, pigs are an ideal species for modeling pediatric brain injuries. All mammalian species experience a transient period of rapid brain growth termed the “brain growth spurt”, and the brain is recognized as being particularly vulnerable to the effects of developmental damage during this time (17, 22). While rodents are postnatal brain developers, and sheep and non-human primates are prenatal brain developers, humans and pigs, being perinatal brain developers, both experience a brain growth spurt near the time of birth (16).

Indeed, the pig has long been considered a highly valuable animal model for biomedical research (23–26). Pigs have frequently been used for research in toxicology (27, 28), diabetes (29, 30), imaging (31, 32), and cardiovascular disease (33, 34). For more than a decade, pigs have also been gaining popularity as a desirable large animal model for neuroscience, cognitive and behavioral research (35–38). Agricultural bred pigs are most commonly used in research due to their wide availability and relative low price. However, these pigs can be particularly challenging to work with as they grow quickly and body weight at maturity can easily reach 300kg (35).

Recently, laboratory purpose-bred miniature pigs (minipigs) are being used more frequently in research studies. Breeds such as the Yucatan and Hanford pigs reach an adult body weight of 70-90 kg, and micropig breeds such as the Gottingen and Sinclair pigs reach an adult weight of 35-55 kg (35). The compact size and the docile temperament of these minipigs make them an attractive model for brain research.

It is conceivable that porcine cognitive and motor performance is dynamic and changes with age, thus it remains necessary to comprehensively characterize minipig behavior over time. We have conducted a battery of behavioral and physiological tests in Yucatan minipigs using established behavioral testing methods, as well as state-of-the-art wearable sensors and artificial intelligence (AI) methods to analyze behavior. We tested their neuropsychological parameters including, executive functions, processing speed, anxiety and learning and memory. We also tested motor and physiological characteristics including gait analysis, circadian rhythms, and daily activity levels. We describe their cognitive, sensorimotor, and physiological function from adolescence (4 months old) through maturity (7 months old) (39). This multi-modal understanding of healthy Yucatan minipig behavior and how their behavior is dynamic throughout development is essential for identifying behavioral and physiological changes induced by brain injury and disease and to test the efficacy of potential therapeutic strategies.

## Results

Neuropsychological screening for executive function, anxiety, willingness to explore a new environment and locomotion were performed using the open field test (40). The open field test has been used in many animal species, including rodents (41–43) cows (44), sheep (45), as well as pigs (46–48). In this study, pigs were placed in a 1.83 m × 1.83 m open chamber and their activity was recorded using overhead cameras for 10 minutes, once per week. We used deep-learning artificial intelligence (AI) software, DeepLabCut (49, 50), to track pig locomotor activity in the open field arena. **Figure 1** shows the distance travelled by pigs (n=4) in the open field from 18 weeks of age to 36 weeks of age. The results show that on average the pigs walked 217± 20 m, but between the age of 27-29 weeks old there was a significant increase in walking (396 ± 19 m). RStudio analysis of coordinate data extracted from DeepLabCut allowed us to calculate the spatial distribution of the pigs’ movement and we found that pigs spend much of their time near the door/entrance to the arena. **Figure 2** characterizes attempts by the pigs to escape the arena defined as any behavior where the pigs pushed on the walls or door of the chamber. Figure 2A illustrates the number of individual events where pigs would exhibit this behavior, whereas 2B shows the cumulative seconds pigs spent engaged in escape behaviors. **Figure 3** shows heatmap representations of pig location within the chamber. Figure 3a is a composite heatmap of all 4 pigs during the open field test across all weeks overlaid with an overhead still image of a pig during the test period. Our results confirm our observations that pigs spend much of the test period in the area immediately surrounding the door (within 30 cm). Further analysis shows that pigs’ spatial distribution differs with age. Figure 3b is a heatmap representation of one pig at 18 weeks of age, whereas 3c shows the heatmap for the same pig at 36 weeks of age. When pigs are younger, they are less inclined to explore the arena. These data are indicative that young pigs may be more anxious than adults.

**Figure 1:**
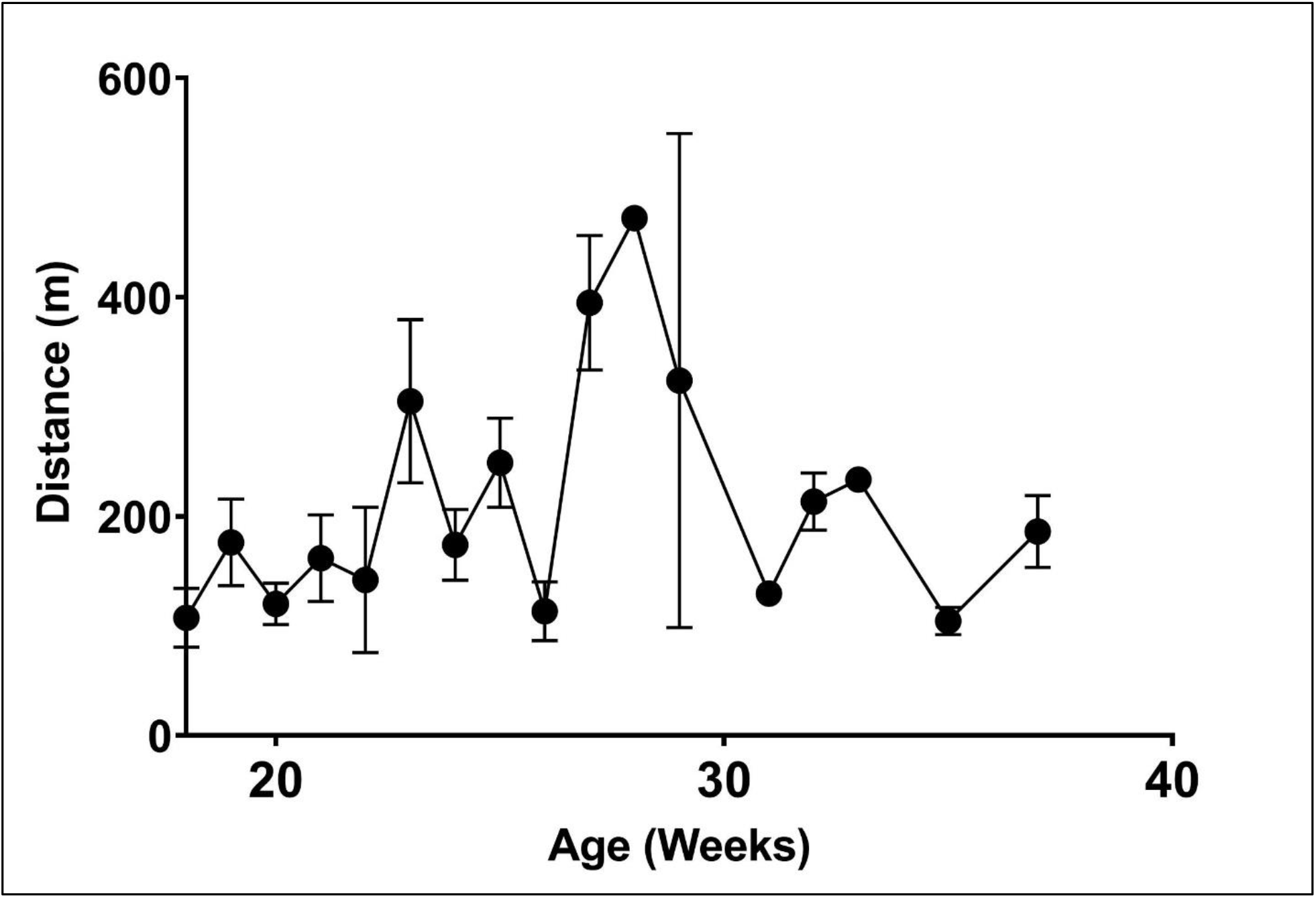
Locomotor Activity in the Open Field. The total average distance minipigs traveled in the open field arena.

**Figure 2:**
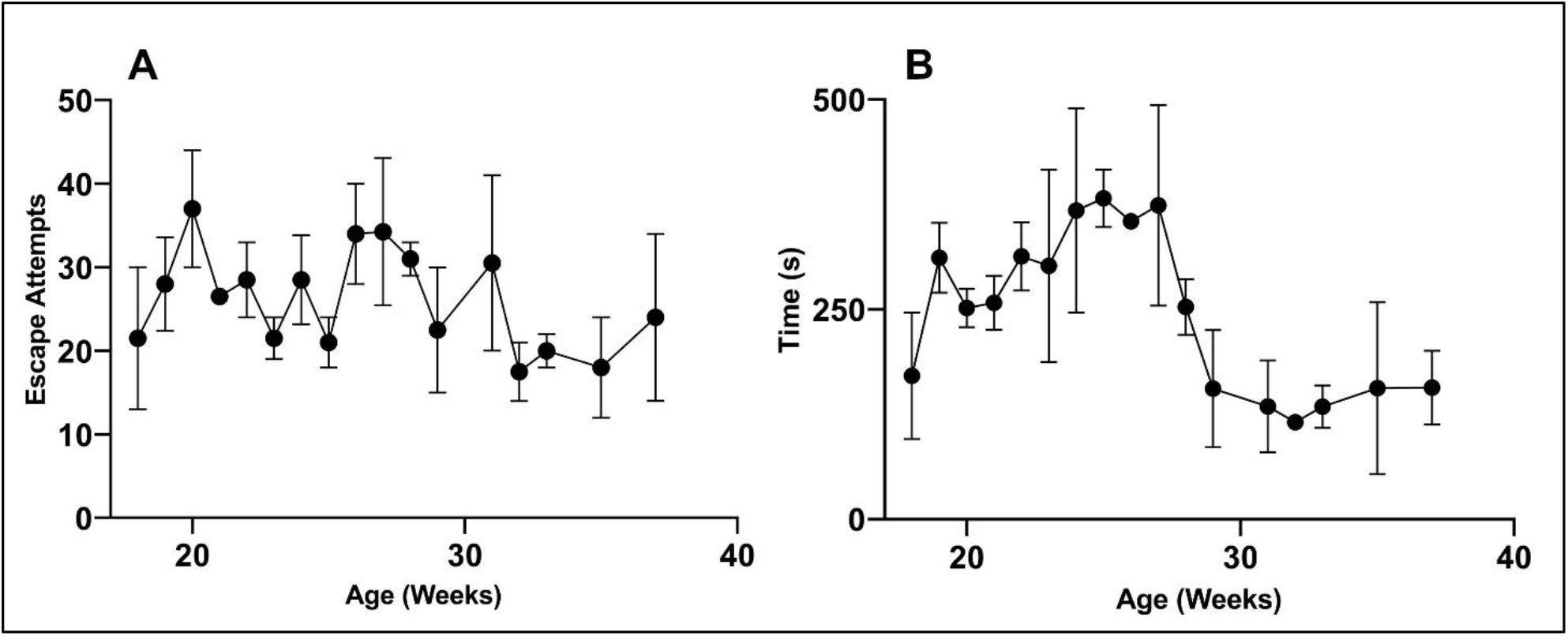
Escape Attempts in the Open Field. A. Number of times pigs actively attempted to escape the arena during the open field test. Escape attempts consisted of pushing on the walls of the arena with the snout or by rearing on the back two legs. B. Total duration of seconds pigs spent actively attempting to escape the arena.

**Figure 3:**
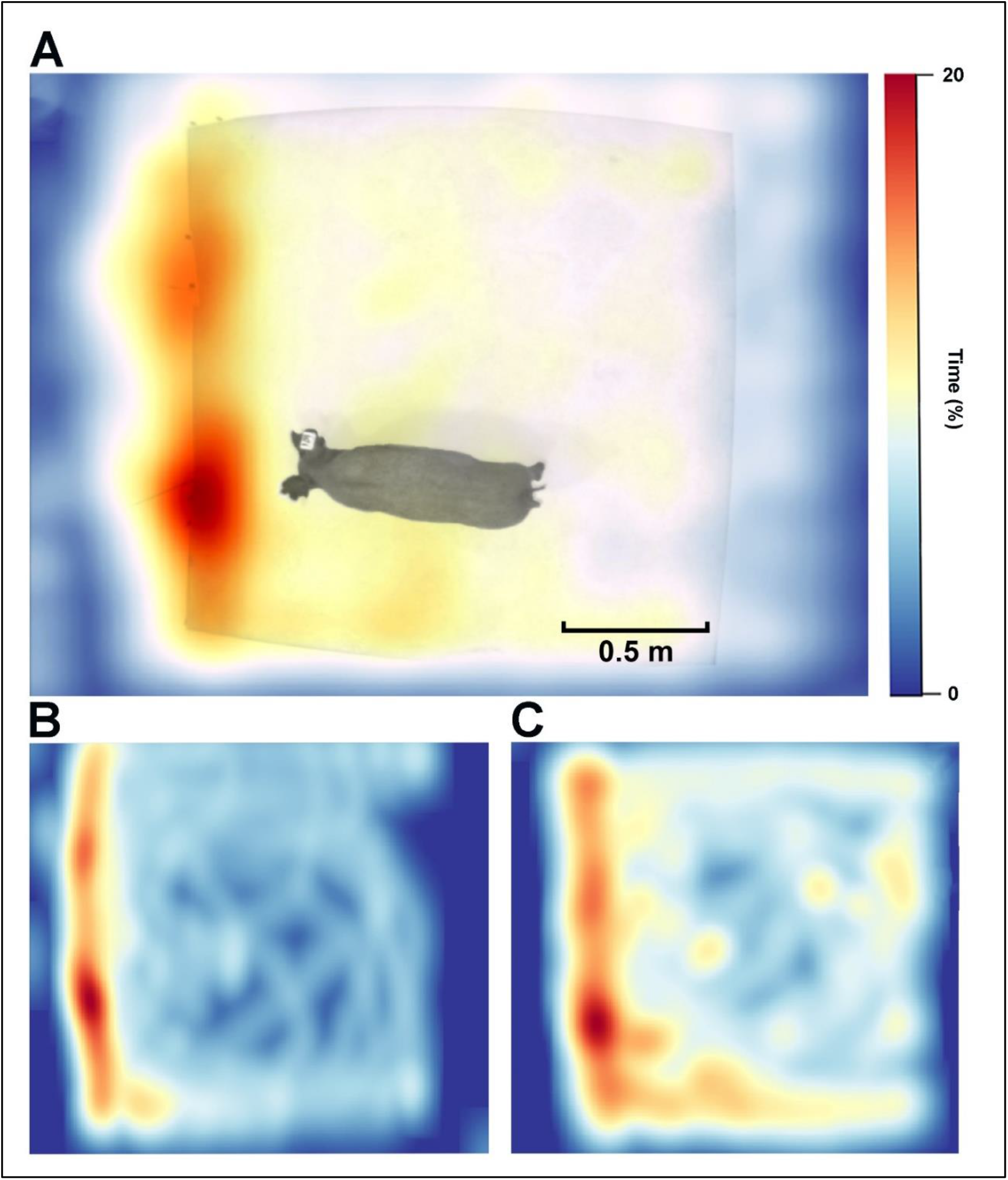
Heatmap representation of pig activity in the open field arena. A. Composite heatmap of pig location from 18-36 weeks. B. Spatial distribution of single pig at 18 weeks of age. C. Spatial distribution of same pig at 36 weeks of age. Healthy pigs exhibit an affinity for the region of the arena closest to the door. This is consistent with findings showing that pigs spend much of their time in the arena attempting to escape.

Measurements of learning and memory, anxiety and depression were performed using the novel object recognition test (51, 52). The novel object task takes place in two parts: the habituation phase and the test phase. Briefly, pigs (n=4) were brought to the behavior chamber and presented with two identical objects during the habituation phase. The pigs were allowed to explore these objects for 10 minutes. The pig was returned to the housing room and after a 15-minute inter-trial interval, the pig was brought back to the behavioral chamber for the test phase. The pig was presented with one of the objects from the habituation phase (familiar object), and one object they had never seen before (novel object). The quantity and duration of contacts with both the familiar and novel objects were recorded. The test took place once per week from 19 weeks to 36 weeks. **Figure 4** shows results from the test phase of the novel object recognition task. Figure 4A illustrates the cumulative duration of contact with the familiar and novel objects. Throughout development, pigs display a significant preference for exploring an unknown, new object (Familiar: 8.2 ± 0.4 s, Novel: 65.1 ± 1.5 s). 4B shows the percent of time the pigs interacted with the novel and familiar objects out of the total time pigs interacted with objects (Familiar: 15.8 ± 0.45 s, Novel: 84.2 ± 0.45 s). These results confirm that healthy pigs prefer the novel object.

**Figure 4:**
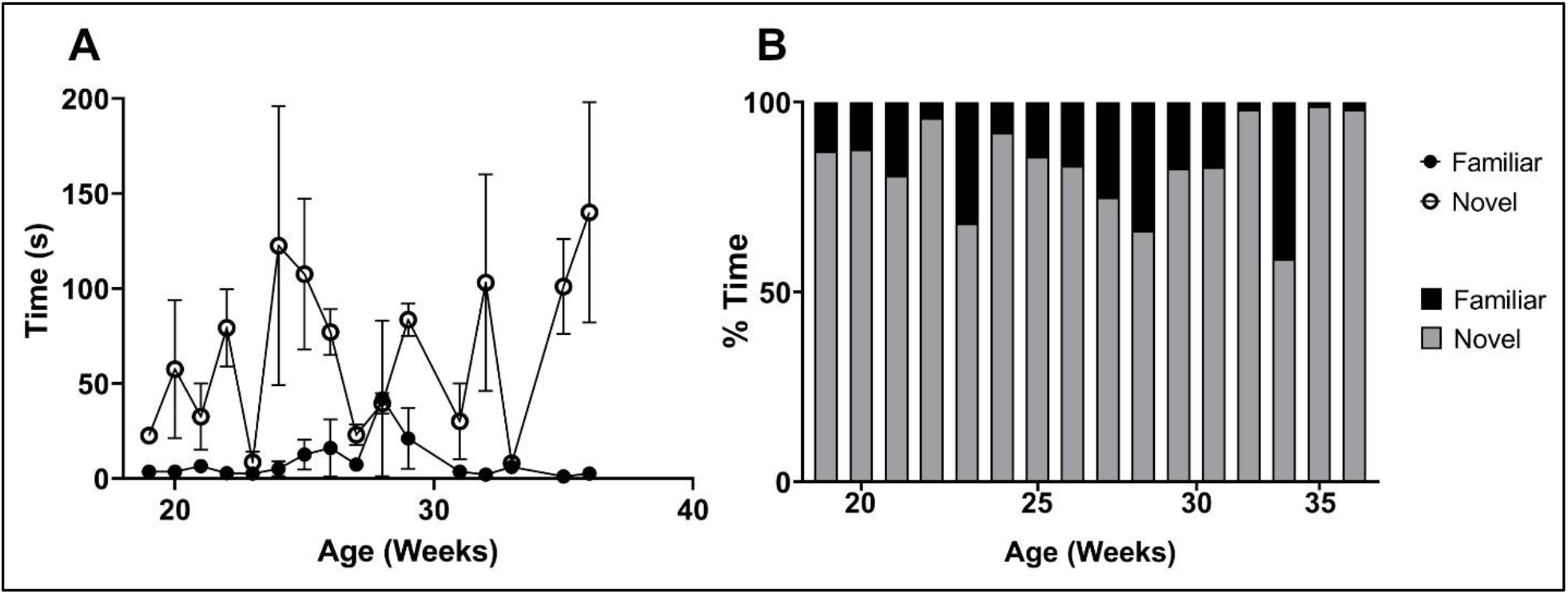
Novel Object Recognition. A. Cumulative duration of contact with objects during the novel object recognition task. Interactions with the familiar object are represented with solid black circles, whereas the interactions with the novel object are represented by open circles. Pigs preferred to interact with a previously unseen novel object than with an object with which they had been previously habituated. B. Average percentage of time pigs spent interacting with objects, split between interactions with the familiar object (black) and the novel object (gray). Pigs consistently interacted with the novel object more than the familiar object.

We have developed a novel test for assessing executive function, processing speed and spatial learning and memory which we term the baited ball pit. This test was inspired by the cognitive hole-board test (53, 54) and is designed to utilize the natural rooting behavior of the pig. In this test, the pig is required to find six apple slices that are hidden in the ball pit. The slices are always hidden at the same location and the time it takes the pig to retrieve each of the six slices is determined. We hypothesize that this test can be used as a measure of situational memory and contextual conditioning. **Figure 5** demonstrates that throughout development the pig became increasingly faster at identifying the hidden objects. On the first week that the test was administered at age 19 weeks, the pigs completed it within 26.5 ± 0.23 s, but increased their speed at finding the object by 100% when they reached 36 weeks old (7.8 ± 0.09 s; n=4).

**Figure 5:**
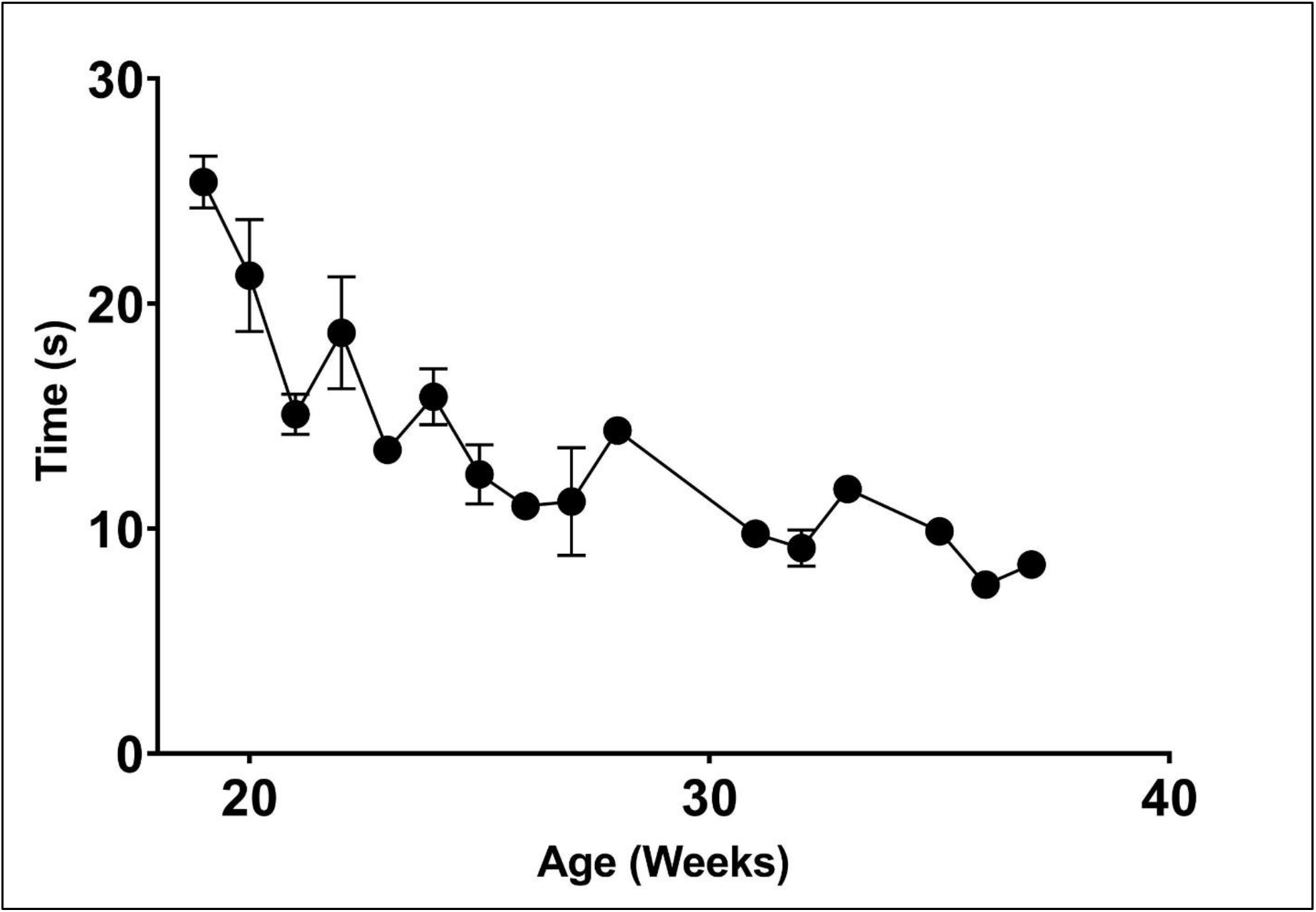
Latency between successful food reward retrievals in the ball pit. Pigs exhibit increased rate of successes with age. As pigs are exposed to the test, they become better at fining apple slices.

The pigs’ natural circadian rhythms were assessed in 5-month-old pigs. We combined results from activity tracker and night-vision video recording to determine the pigs wake and sleep cycles. **Figure 6** shows that pigs are most active between the hours of 12 pm and 4 pm. Pigs are fed at 8am and 3pm. **Figure 7** shows that pigs have consistent sleep cycles, waking around 7 am and sleeping around 11 pm, sleeping for an average of 8.7 hours (± 0.2 h).

**Figure 6:**
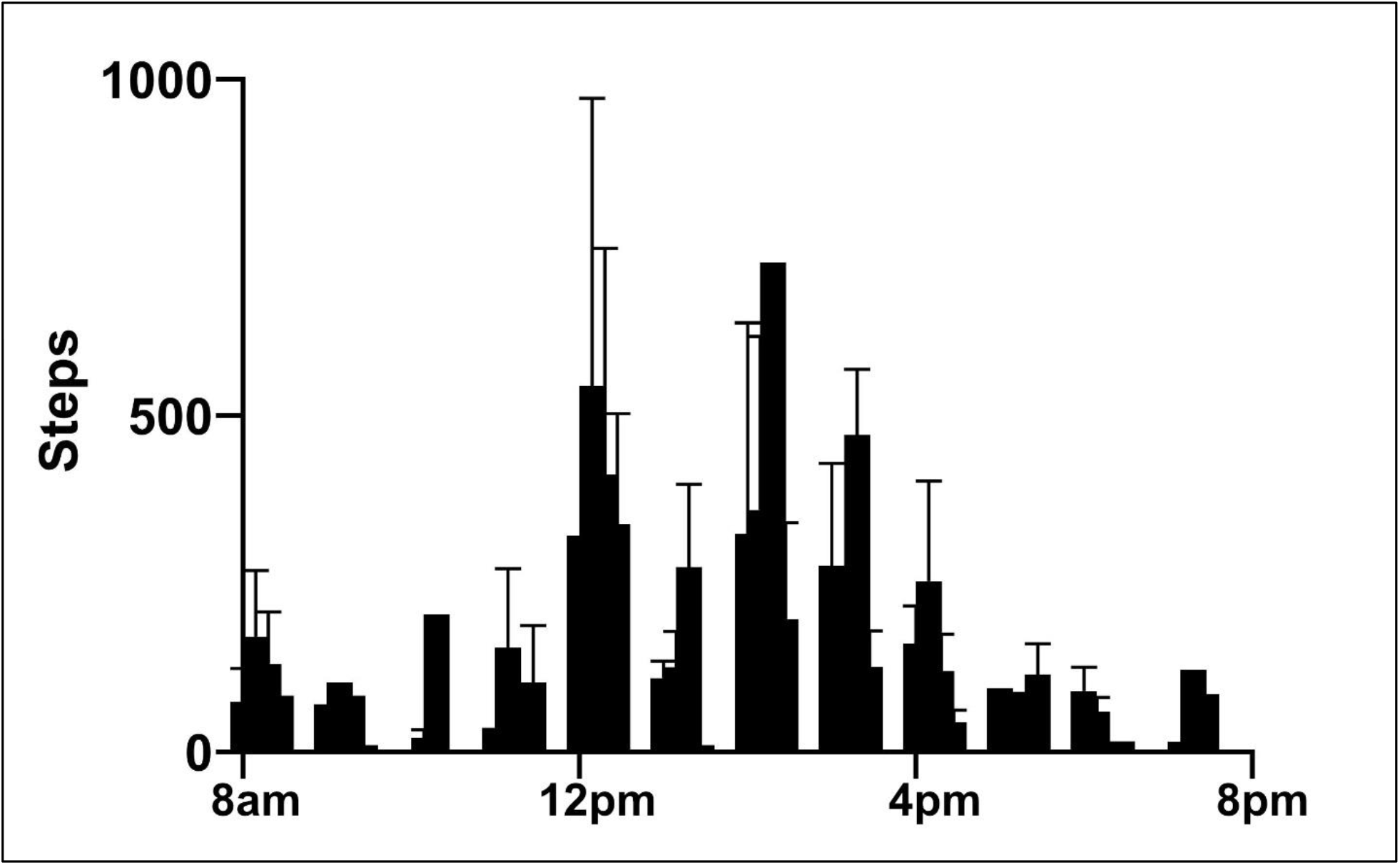
Activity tracking/Fitbit step counting. 12-hour (8:00am-8:00pm) graph showing steps recorded by an activity tracker (Fitbit) on pigs averaged over 5 days, broken into 15-minute bins. Pigs were most active in the middle of the day, between noon and 4pm.

**Figure 7:**
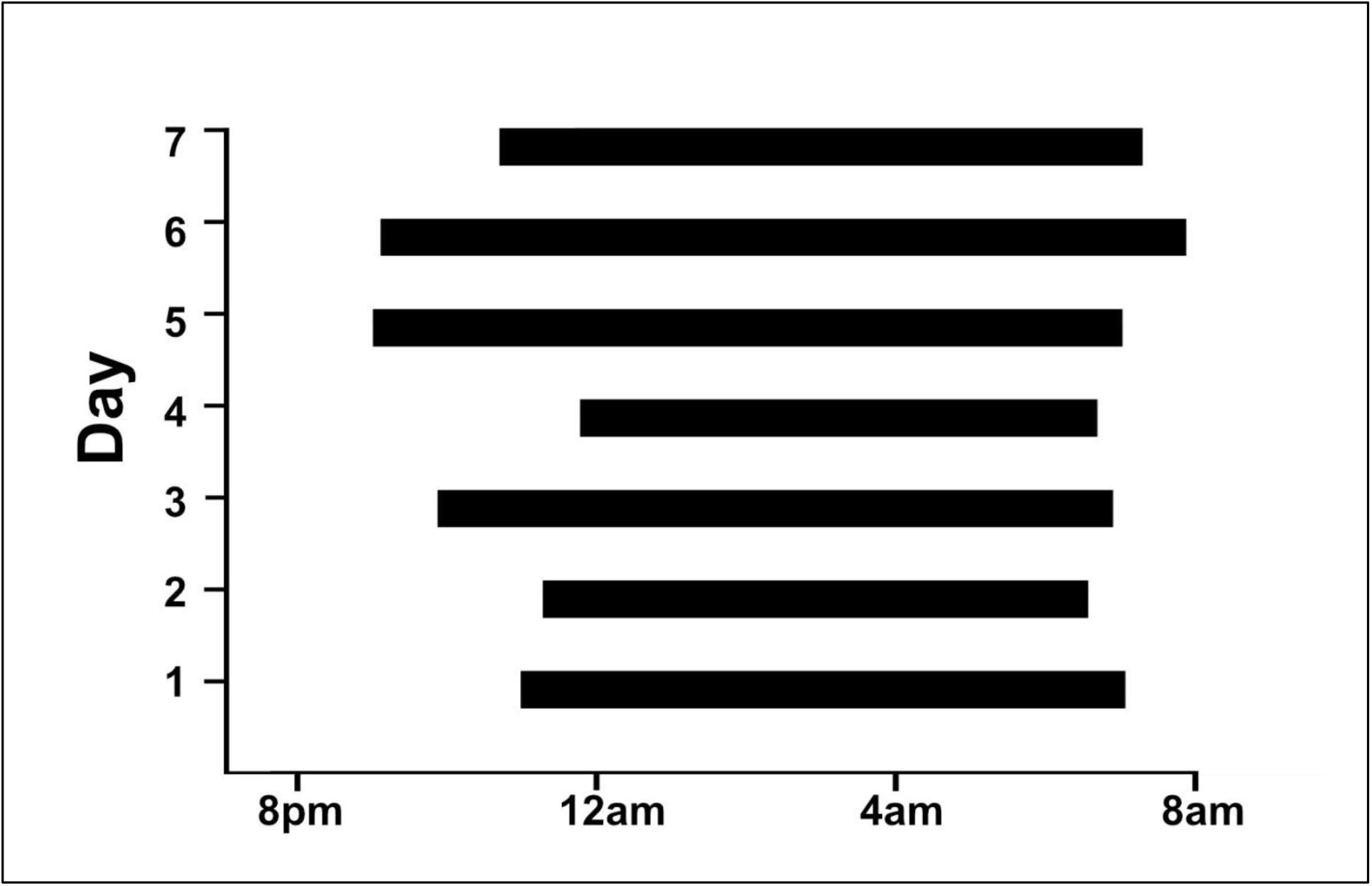
Sleep and Circadian Rhythms. Sleep-Wake graph showing that pigs tend to fall asleep between 10 pm and 12 am and tend to wake around 7 am. Pigs slept for an average of 8.7 hours per night.

We also acquired high resolution kinematics and gait analysis. Three-dimensional (3D) kinematic data has the ability to capture unique parameters that are unable to be identified through video alone. Additionally, location and movements of anatomical points can be quantified with high levels of accuracy through a robust calibration and use of a multi-camera system. Thus, for this study, motion capture was used to obtain initial 3D movement profiles of a pig during a walking trial. As proof of concept, initial kinematic data were calculated for a single pig. The camera space permitted capture of one full gait cycle. One gait cycle is defined as the following touch point in sequence: left hind limb to left front limb followed by right hindlimb touch and right front limb (55).

Three rigid marker pods were placed on the spine of the pig. Each pod consisted of three reflective markers with one located on the head (covering the base of the skull to the second cervical vertebra), a second on the shoulders (directly over the third thoracic vertebrae), and a third on the rear (directly over the sixth lumbar vertebrae). Additionally, four single markers were placed on the front and hind limbs distal to the condyle of the humerus and distal to the lateral condyle of the femur, respectively.

Initial kinematic analyses of the pig gait trials were performed. This initial analysis included the average linear velocity, along with angle data. The angle data were rotations computed about three axes that were created at each pod. For example, an axis perpendicular to the base of the pod was used to define left/right rotation of the head. An anterior axis along the base of the pod was developed and rotation about that axis was left/right lateral tilt and finally a medial/lateral axis was used to define the up/down – or “nodding” motion. These same three axes were computed on the other two pods as well. **Figure 8** shows a sample of the head movement from this pig relative to the global coordinate system.

**Figure 8.**
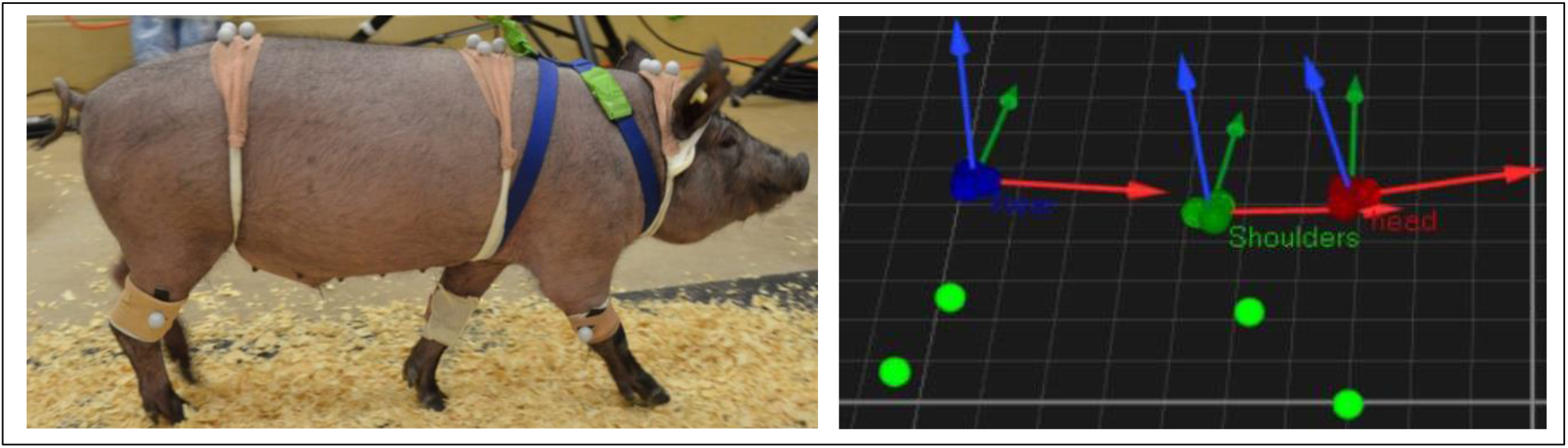
Pig with reflective markers. Three rigid pods are located along the spine (labeled: head, shoulders, and rear) with four single markers, one on each leg.

Data were analyzed of each pod with respect to the fixed reference system (i.e. global coordinate system) The average linear velocity of this pig was 1.93 m/s. Data from this pig showed lateral tilt ranges around 12°. Additionally, the head flexion-extension or “nodding” had a range of approximately 17°. Left to right rotation of the head ranged for this trial showed a 30° movement to one side as the pig moved its head in an attempt to turn. Similar motions were obtained for each of the marker pods located on the shoulders and buttocks regions.

## Discussion

The minipig possess many physiological and anatomical attributes that are similar to humans, making it an attractive animal model for biomedical research and an emerging translational model in neurology and neuroscience. The ability to produce in the minipig brain an injury of similar nature to humans (2, 56–58), and the opportunity to acquire neuropsychological and motor performance measurements, make this animal model invaluable in the field of neurology and rehabilitation. It is of particular interest to understand how different features of the behavior change throughout age. In adults, recovery after brain injury is considered when performance returns to pre-morbid levels, but in children and adolescents this is more complex. During the time that it may require for children who suffered brain injury to return to pre-morbid levels, other children may have reached new milestones and the critical period of acquiring or improving these abilities may have passed, leading to long term impairments. Therefore, characterizing the dynamic of cognitive, sensorimotor, and physiological functions in the minipig model is critical for translational research.

We have used a battery of neuropsychological tests to evaluate processing speed, learning and memory, and anxiety in the minipigs. These tests are particularly relevant for assessing the severity of brain injury in animal models (42, 59–65) and are highly translational in human research. Similar to humas, the minipigs show increased learning capabilities, long-term memory, and increased self-confidence with age.

The physiological measures are also relevant for neurology and rehabilitation research. Brain injury is often accompanied by reduced quality of sleep and sleep disturbances (66–69). The results demonstrate that a combination of activity tracker and video recording can provide valuable data regarding these aspects of behavior.

We have used AI to analyze 400 minutes of video recording, that would have taken as much time, or more, to be manually analyzed. Machine learning and AI methods are becoming increasingly useful in behavioral research and provide new opportunity to analyze large sets of data with high temporal and spatial resolution and in less time than it would have taken by a manual user.

This work also demonstrates the initial data sets possible with a 3D kinematic gait analysis. In the future we will take rotations of each pod location (head, shoulders, rear) and compare the motions between each whereas the above data are the movement of the pods with respect to a global system. Future work will use this approach to quantify and compare changes in movement pre and post trauma within and across pigs. We will gather data on several healthy pigs to obtain a profile of healthy motions. Additionally, pigs with injuries or diseases could be tested to identify how gait parameters are affected by a specific disease or injury.

While there exists an ample amount of literature describing pig anatomy and physiology, studies investigating the complexity of pig psychology and behavior are less abundant (38, 46, 70–72). Much of this scarcity can likely be attributed to the numerous challenges associated with the use of large animals in research. Due to handling concerns, agriculture pigs are typically used only for acute studies, with the age cap at 5-6 weeks of age (35). This tends to prevent long-term behavioral studies from being done.

This study extends the toolbox of behavioral and physiological tests in the Yucatan minipig. These comprehensive measurements of different aspects of behavior can be useful to measure performance after major and mild injuries and may be sensitive to subtle changes in performance that are often difficult to diagnose clinically.

## Methods

### Animals

All experiments were conducted in compliance with guidelines set by the Michigan State University Institutional Animal Care and Use Committee.

Four Yucatan minipigs, 2 female 2 male, were used in this study (Premier BioSource, CA). Behavioral testing began when pigs were 4-months of age. Pigs were pair housed in an enriched environment, fed nutritionally complete feed twice per day, with unrestricted access to water on a 12-hour (7:00-19:00) light cycle.

### Behavioral Chamber

Behavioral experiments, with the exception of gait analysis, took place in an open chamber measuring 1.83m × 1.83m. The walls of the chamber were made of commercially available PVC board and measured 1m in height. A bullet camera (Omron Sentech) was suspended overhead to record locomotion. The concrete floor of the chamber was hosed down between subjects.

### Behavioral Experiments

#### Open Field

Pigs were individually led to the behavioral chamber. Each pig was allowed to freely explore the chamber for 10 minutes and locomotor activity was recorded by the camera suspended overhead. This test was conducted once per week.

#### Novel Object

Pigs were individually led to the behavioral chamber. Pigs were first exposed to two identical toys and all activity was recorded by the camera suspended overhead. This initial habituation phase lasted 10 minutes. The pig was then returned to the housing room for a 15-minute inter-trial interval. Following this interval, the pig was again led to the behavioral chamber and presented with one familiar object from the habituation phase and one new/novel object that had not been seen before. The test phase lasted 10 minutes, after which the pig was returned to the housing room.

#### Ball Pit

Prior to testing, handlers placed a plastic wading pool (91.44 × 91.44 × 17.53 cm) in the behavioral chamber and filled it with colorful plastic balls (5.59 cm diameter). Six slices of apple were buried in a pentagonal shape around the perimeter of the pool with one apple slice in the center. Pigs were then individually led to the behavioral chamber. Cameras, both overhead and handheld, recorded as the pigs searched for and retrieved apple slices. The duration of this test was less than 5 minutes.

Analysis was conducted by manually watching the videos and tracking the timestamp at which each apple slice was retrieved. The latency between successful retrievals was calculated, and a mean latency for each trial was determined.

#### Gait Analysis

Pigs were trained to tolerate wearing commercially available miniature swine harnesses and to walk on a lead with the handler.

#### Activity Tracking

Fitness tracking devices (Fitbit) were attached to miniature swine harnesses. Pigs wore harnesses and fitness trackers for several hours per week. Step data was extracted from the Fitbit app.

#### Circadian Rhythms

Wide angle cameras (amazon cloud cam, wyze cam) were used to record pigs in the housing room 24/7. Times that the pigs wake in the morning and fall asleep in the evening were recorded.

## Acknowledgment

G.P. is partially supported by National Institutes of Health grants R01NS072171, R01NS098231 and UF1NS115817.

